# Generating Novel Leads for Drug Discovery using LLMs with Logical Feedback

**DOI:** 10.1101/2023.09.14.557698

**Authors:** Shreyas Bhat Brahmavar, Ashwin Srinivasan, Tirtharaj Dash, Sowmya R Krishnan, Lovekesh Vig, Arijit Roy, Raviprasad Aduri

## Abstract

Large Language Models (LLMs) can be used as repositories of biological and chemical information to generate pharmacological lead compounds. However, for LLMs to focus on specific drug targets typically require experimentation with progressively more refined prompts. Results thus become dependent not just on what is known about the target, but also on what is known about the prompt-engineering. In this paper, we separate the prompt into domain-constraints that can be written in a standard logical form, and a simple text-based query. We investigate whether LLMs can be guided, not by refining prompts manually, but by refining the the logical component automatically, keeping the query unchanged. We describe an iterative procedure LMLF (“Language Models with Logical Feedback”) in which the constraints are progressively refined using a logical notion of generalisation. On any iteration, newly generated instances are verified against the constraint, providing “logical-feedback” for the next iteration’s refinement of the constraints. We evaluate LMLF using two well-known targets (inhibition of the Janus Kinase 2; and Dopamine Receptor D2); and two different LLMs (GPT-3 and PaLM). We show that LMLF, starting with the same logical constraints and query text, can guide both LLMs to generate potential leads. We find: (a) Binding affinities of LMLF-generated molecules are skewed towards higher binding affinities than those from existing baselines; LMLF results in generating molecules that are skewed towards higher binding affinities than without logical feedback; (c) Assessment by a computational chemist suggests that LMLF generated compounds may be novel inhibitors. These findings suggest that LLMs with logical feedback may provide a mechanism for generating new leads without requiring the domain-specialist to acquire sophisticated skills in prompt-engineering.

## 1 Introduction

In 1982, Edward Feigenbaum identified a difficulty in the transfer of human-knowledge to a machine, now famous as “the Feigenbaum bottleneck” (Feigenbaum et al. 1977). In a curious twist of fate, we now appear confronted by a “reverse bottleneck”. Machine knowledge, such as those contained in large foundation models, is at least as difficult for humans to access as it was to represent human knowledge in a machine-understandable form. Surprisingly, this reverse bottleneck also appears to have been first identified in 1982. The problem of ‘The Human Window’ (Kopec 1982; Michie 1982) refers to the difficulties faced by humans when interacting with a complex computing system, due to a mismatch in representations of the human and machine. Modern large language models (LLMs) would appear to have resolved this difficulty through their impressive facility to use natural language as a mechanism of communicating with humans (Brown et al. 2020; Narang and Chowdhery 2022). In fact, the true difficulties concerning the Human Window arise from the need for a *conceptual* interface, not simply a linguistic interface. That is, the mismatch has to be addressed at a concept-level.^1^ The intense interest in the methods and practices of ‘prompt engineering’ as an approach to extract useful information from LLMs could be seen as evidence of the deeper, conceptual mismatch that exists between LLMs and human representations. In this paper, we are concerned with an immediate practical manifestation of this, namely in the apparent need to be a sophisticated prompt-engineer in order to be able to use the capability of an LLM best.

Our specific interest is in the generation of novel leads for early-stage drug-design. Here, the human is typically a computational-or synthetic-chemist, who often uses knowledge that can be expressed in a logical form. For example, this may be in the form of generic constraints on values like molecular weight, hydrophobicity, synthetic accessibility score, *etc*.; and specific constraints like estimated binding energy to the target site, size of the binding site, presence of any specific anchors and so on. The usual route to provide this as input to an LLM would be through prompts, which combine what the chemist knows, and what the chemist needs from the LLM. However, the free-text interface prompts make it difficult to settle on a single form for this input, and the process becomes one of experimentation with phrasings or text, and orderings of textual sequences. Results are, therefore, dependent not just on what is known bio-chemically, but on the content and sequence of text provided. This makes the experiments highly subjective and difficult to reproduce.

In this paper, we attempt to reduce this subjectivity, while attempting to retain one very important feature of LLMs, namely, the ability to adapt quickly to new probability distributions and to sample effectively from them. Our approach is to separate the content of a prompt into two parts: a domain-specific component, and a domain-independent component. In this paper, by ‘domain-specific’ here, we mean the drug-design specific drug-design aspects that allow the LLM to update its probability distribution. The domain-independent part will be a simple query enabling the LLM to generate instances from its (updated) distribution. Further, we will require that the domain-specific component can be encoded in a standardised form that can be refined automatically; and the domain-independent form is simple enough to be independent of the LLM used. This is done to ensure clarity and repeatability of the experimental protocol. Here the standardised form we employ is formal logic, and LLM updates are done through a procedure LMLF that employs what we call ‘logical feedback’.

The rest of the paper is organised as follows. In Sec. 2 we summarise the need for automated discovery of new leads in early-stage drug design, and a description of some constraints on lead-generation. Although we expect readers to be familiar with LLMs, at least in their use through interactive interfaces like ChatGPT, we nevertheless include a short description of LLMs as implementations of complex probabilistic generative models. Section 3 we describe the LMLF procedure as a method of using LLMs in conjunction with satisfaction of logical constraints acting as feedback to update its probability distribution. Section 4 evaluates LMLF empirically using two benchmark drug-design targets and two well-known LLMs. Related work is in Sec. 5. Section 6 concludes the paper.

## 2 Background

### 2.1 Lead Discovery in Early-Stage Drug Design

Drugs are small molecules that usually attach themselves to parts of larger molecules (like proteins). This attachment takes place at a location known as the ‘target site,’ and the larger molecule is itself sometimes referred to as ‘the target.’ The attachment occurs mainly by the usual physical electrostatic mechanisms, and the process is known as *binding* The target molecule with the small molecule bound to it has a different shape from the original molecule. It is generally thought that function depends on the shape, and therefore the functioning of the large molecule is changed. Usually, this change means stopping some activity, and the small molecule is said to *inhibit* the target. Leads are small molecules that could potentially bind to a target molecule.

Artificial Intelligence is currently revolutionizing drug development, especially in various steps of early-stage drugdesign (see Fig. 1(a)) as virtual screening, identifying qualitative and quantitative structure-activity relations (SARs) and so on. The broader picture is of a semi-automated scientific discovery pipeline involving feedback from from computational chemists, synthetic chemists and biologists and manufacturers, using results from simulation, synthesis protocols, and biological testing (see Fig. 1(b) and (Zenil et al. 2023) for the broader context of closed-loop scientific discovery).

**Figure 1.**
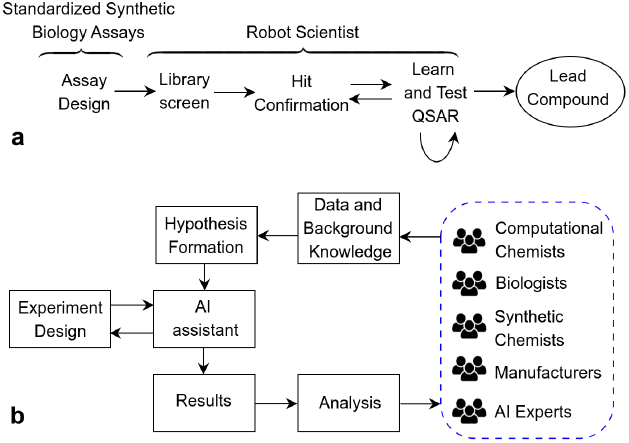
(a) Early Stage Drug Design; and (b) Computational drug discovery with specialists-in-the-loop.

In this paper, we restrict ourselves to a single domain-specialist in the form of a computational chemist with a knowledge of chemical synthesis. We envisage 2 kinds of interaction between such this specialist and the computational engine: (A) Provision of chemical knowledge. This could be of a general nature on drug-likeness, or specific to the target or output of the computational engine; (B) Asking chemical queries, usually about possible new structures, or specific aspects of existing structures.

If the computational engine is a large language model (LLM), then all specialists in Fig. 1(b) should be able to interact using natural language. But, as pointed out in Sec. 1, the very flexibility allowed by natural language instruction poses difficulties to the construction a pipeline capable of repeatable, standardised experiments. We will be looking at a mechanism that requires: the specialist’s knowledge (A) to be provided in a standardised form that can then be refined automatically; and the chemical queries (B) that are to be posed as simple text concerning the generation of new molecular structures. In effect, (A) and its subsequent refinements are used to alter automatically the conditioning information (used here in a probabilistic sense) provided to the LLM; and (B) is used to sample from the resulting conditional probability distribution over small molecules.

### 2.2 Language Models as Probabilistic Machines

A language model is a probabilistic model of natural language that learns a probability distribution over sequences (or sentences) of tokens. Let *W* denote one such sentence with *N* tokens, *W* = (*w*_1_, …, *w*_*N*_). In practice, *N* is arbitrary. So, a language model estimates the probability of observing the sentence (*w*_1_, …, *w*_*N*_), denoted by the joint probability *P* (*w*_1_, …, *w*_*N*_). Mathematically, this joint probability can be expanded using the chain-rule of probability as

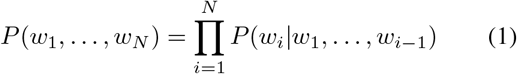

In practice *P* (*w*_1_, …, *w*_*N*_) is approximated using *n*-gram models (Jelinek 1980; Katz 1987) or Neural language models or NLMs (Bengio, Ducharme, and Vincent 2000). These models make Markov assumption that the probability of a word depends only on the previous *n* < *N* words. That is,

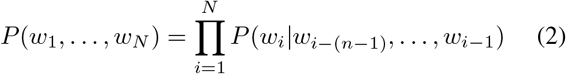

Neural language models or NLM (Bengio, Ducharme, and Vincent 2000) are probabilistic language models based on (deep) neural networks that can handle the problems associated with *n*-gram models, such as handling long-range dependency, context understanding, handling noise and ambiguity, learning complex relationships by learning a distributed representation of text tokens. An NLM approximates each term on the right-hand side of Eq. (2) using a neural network. Large Language Models (LLMs) such as GPTs (Radford et al. 2019) and PALM (Narang and Chowdhery 2022) are large and complex neural language models that use the transformer architecture (Vaswani et al. 2017) to learn from vast and diverse corpora of text data.

Prompt-based LLMs, such as ChatGPT and BARD are LLMs that can learn with human-feedback (Ouyang et al. 2022). A prompt is an input sequence, written by a human in a natural language serves as the starting point that sets the context of the LLM to generate the next (highly) probable text sequence in an auto-regressive manner.

## 3 Using Language Models with Logical Feedback (LMLF)

We differentiate the information provided by a human in a prompt for an LLM into 2 parts: contextual information, which can be encoded in a formal language, and a query, which is in a natural language. It is further helpful for us to distinguish the former into background knowledge *B* consisting of definitions, functions, procedures and factual statements, and *C*, consisting of domain-constraints. For the specific task of lead-discovery that we are interested in:

- *B* will include: example molecules; facts about the molecules obtained from computation by a general-purpose molecular modelling package (computing, for example, bulk properties like molecular weight, synthesis accessibility scores *etc*.); facts about the molecule obtained from computation by special-purpose molecular modelling package (computing for example, binding affinity to the target site). We also consider part of *B* any standard mathematical and arithmetic procedures used in the constraints *C*.
- *C* will typically be a conjunction of desirable properties of leads, like molecular weight between 200 and 700; logP below 5; SA score below 5; and binding affinity is −8 or less, etc.

In developing LMLF, we are inspired by the MIMIC algorithm (De Bonet, Isbell, and Viola 1996), which uses an iterative procedure for model-assisted sampling. MIMIC assumes that we are looking to generate instances with low values of an objective function *F*. On any iteration *i*, MIMIC has access to a sample of instances; and a model *M*_*i*_ that can be used both for discrimination and for sampling (generation). True *F* -values are computed for the sample of known instances, and *M*_*i*_ is revised to *M*_*i*+1_ that can discriminate accurately between instances with *F* -value below and above some threshold *θ*_*i*_. That is, *M* discriminates between *F* (*x*) ≤ *θ* (labelled “1”) and *F* (*x*) *> θ* (labelled “0”). *M*_*i*+1_ is then used to generate new data instances, the threshold *θ*_*i*_ is lowered to *θ*_*i*+1_, and the process is repeated (say *k* times). We first recast MIMIC in terms of background knowledge and constraints. Let *B* denote the function *F*, the thresholds *θ*_*i*_, and standard arithmetic definitions of ≤, *>*. Let *C* be the conjunction *C*_1_ ∧ *C*_2_ ∧ ⋯ *C*_*n*_, where *C*_*i*_ = (*F* (*x*) ≤ *θ*_*i*_). we are able to abstract two general principles about the algorithm:

- On any iteration *i*, feedback to *M*_*i*_ is provided by instances labelled based on whether (*B* ∧ *C*_*i*_) is true (label 1) or false (label 0). We call this the “constraint-based labelling” property.
- Since *θ*_*i*+1_ ≤ *θ*_*i*_, if *F* (*x*) ≤ *θ*_*i*+1_ then *F* (*x*) ≤ *θ*_*i*_. That is, (*B* ∧ *C*_*i*+1_) |= (*B* ∧ *C*_*i*_). We call this the “constraint-generalisation” property.

We now devise a general iterative procedure with these two properties to alter the conditioning sequence for an LLM for discrimination and generation. The steps are shown in Procedure 1. For reasons of space, we do not provide procedures for the auxiliary functions. An idealised worked example below is intended to help clarify their intended behaviour.

### Procedure 1: Incremental sampling from an LLM’s conditional distribution using iterative constraint-based labelling and constraint generalisation.

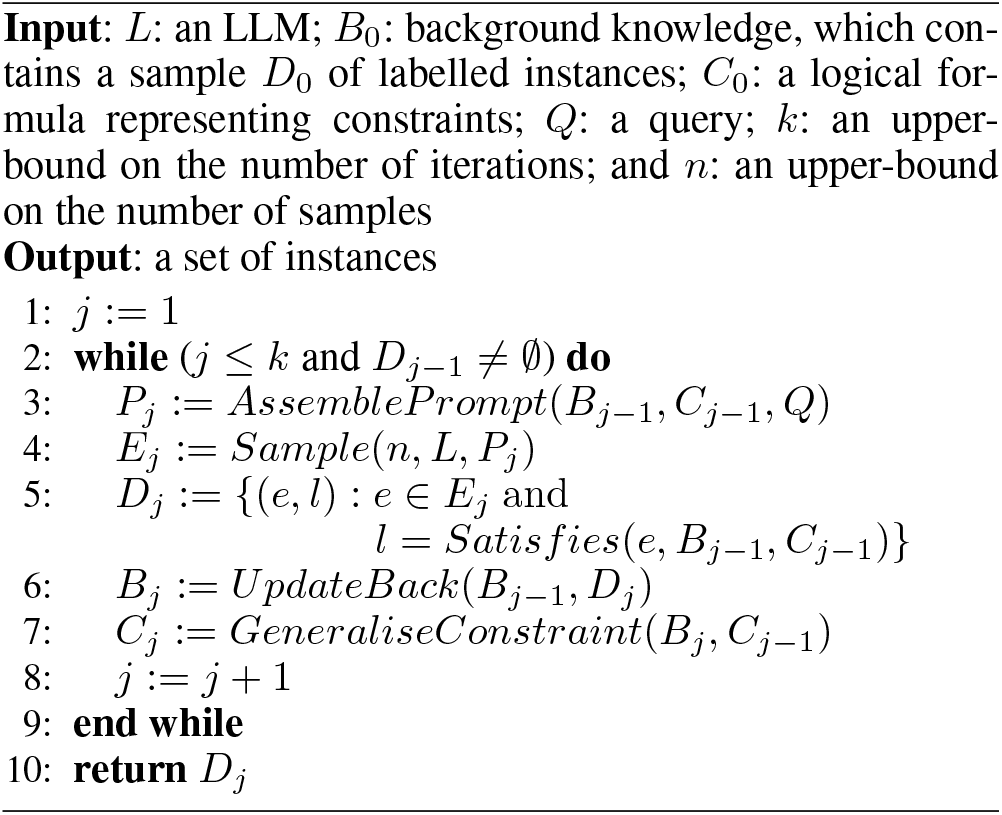

**Example 1**. *We want to generate molecules to inhibit the target protein, Janus Kinase 2* (*JAK2*). *We work through one complete iteration of LMLF, albeit without actual details. For ease of explanation, we will be using a logic-based syntax to describe background knowledge and constraints, and restricted natural language* (*Kuhn 2014*) *to describe the query* (*the actual implementation used in experiments does not use either of these representations*).

1. *The background knowledge B*_0_ *contains facts about the target, some known molecules, and labels* (*say, 1 and 0 inhibitors*).
  - *These are conjunctions of facts. For example: target*(*jak*2) ∧ *mol*(*m*_1_) ∧ *label*(*m*_1_, 1) ∧ *mol*(*m*_2_) ∧ *label*(*m*2, 0) ∧ ….
  - *Additional facts may describe the properties of the molecules; for example, molwt*(*m*_1_, 245.5) ∧ *logp*(*m*_1_, 4.0) ∧ …. *B*_0_ *also contains information for use by the constraints*.
  - *This may be facts. For example, a threshold for binding affinity, like affthresh*(8.0); *or a range of allowed values for molecular weight, like molwt*([200, 700]) *and so on*.
  - *B*_0_ *also contains functions for performing computation. For example, the definition for computing affinity scores: affinity*(*Mol, Target*) = *Score*) if (`GNINA` (*Mol, Target*) = *Score*), ….
2. *C*_0_ *describes a set of constraints required to be satisfied. Some examples are:*
  - *mol*(*m*_10_) ∧ *molwt*(*m*_10_, *w*) ∧ *molwt*([*x, y*]) ∧ (*x* ≤ *w*) ∧ (*w* ≤ *y*)
  - *mol*(*m*_10_)∧*affinity*(*m*_10_, *a*)∧*affthresh*(*z*)∧(*a > z*). *Here a, w, x, y, z are all variables*.
3. *The query Q we use is a simple textual one: Generate valid SMILES string for n molecules that are labelled “1” and are not found in any known database*.
4. *Using the information in B*_0_ *and C*_0_, *and Q, AssemblePrompt returns text string that includes strings for labelled molecules* (*like* 1 *m*_1_, 0 *m*_2_ *and so on*) *and the query Q. The LLM uses this as a prompt to sample n new molecules*.
5. *Each new molecule e is tested against the constraints. The function Satisfies returns* 1 *if e satisfies B*_0_ ∧ *C*_0_ *and* 0 *otherwise*.
6. *The background knowledge is updated to B*_1_ *with the newly labelled instances*.
7. *The constraint C*_0_ *is generalised to C*_1_ *by GeneraliseConstraint. Generalisation is restricted to numeric constraints with inequalities* (*all other constraints are left unchanged*). *A constraint of the form x* ≤ *θ* (*where θ is some numeric value*) *is generalised to x* ≤ *θ*^*′*^ *where θ*^*′*^ < *θ. Similarly x > θ is changed to x > θ*. *It is assumed that the background knowledge contains a function to compute θ*^*′*^ *given θ* (*for example, increment and decrement functions that add or subtracts pre-specified amounts to θ*).

The implementation used for experiments in the paper has aspects related to efficiency and book-keeping which introduce unnecessary detail, but retains the essential feature of iteration over constraint-based labelling and constraint-generalisation. In the following, we will call the implementation `PyLMLF`, and refer the reader to the code accompanying the paper for its content. For our purpose, `PyLMLF` will be used as a tool for investigating the use of LMLF for the generation of new leads for early-stage drug-design.

## 4 Empirical Evaluation

### 4.1 Aims

We use `PyLMLF` as a tool to investigate the following conjecture:

- The use of LLMs with logical feedback generates better lead molecules for early-stage drug-design than LLMs without logical feedback.

We will make the following design choices to conduct the experiments: (a) We consider two classic drug-design targets and two well-known LLMs; (b) We will use two methods of assessing the results: quantitatively, using the distribution of binding affinities of generated molecules; and qualitatively, using assessments by a computational chemist.

### 4.2 Materials

#### Biological Targets and LLMs

We conduct our evaluations on JAK2, with 4100 molecules provided with labels (3700 active) and DRD2 (4070 molecules with labels, of which 3670 are active). These datasets were collected from ChEMBL (Gaulton et al. 2012) which are selected based on their *IC*_50_ values and docking scores with active JAK2 and DRD2 proteins less than −7.0. For all our experiments, we use 2 LLMs: GPT-3.0 (Brown et al. 2020) and PaLM (Narang and Chowdhery 2022).

#### Background knowledge

There are 3 categories of background knowledge: (a) Factual statements: referring to what is already known about drug targets, for example, some subset of experimentally known inhibitors for JAK2 and DRD2; (b) Functional definitions to compute bulk molecular properties; and and (c) Procedures needed to assemble the prompt for sampling. For (b), we use the definitions available within the molecular modelling packages RDKit (Landrum et al. 2013) and GNINA 1.0 (McNutt et al. 2021) to compute the validity of SMILES string, molecular weight, synthetic accessibility score (SAS), LogP, binding affinity. For (c), we use Python’s `f` -string syntax to incorporate the molecules and their class-labels (*inhibitor* or *non-inhibitor*) represented in (a).

#### Constraints

We conduct experiments with 3 categories of constraints: (a) No constraints; (b) Target-agnostic constraints; and (c) Target-specific constraints. Of these, (a) is self-explanatory. Constraints in category (b) refer only to some generic bulk properties of the small molecules. Specifically, we use the generic constraints used in (Dash et al. 2021) for one of the datasets used here (JAK2). These constraints encode the following requirements of potential leads: (i) Molecular weight must be between 200 and 700; (ii) Synthetic accessibility score (SAS) must be below 5; (iii) LogP value must be below 5; and (iv) Binding affinity must be above 7. For experiments here, we limit constraints in (c) to estimates of binding affinity to the target site obtained from the approach described in (McNutt et al. 2021). For a small molecule *m*, the constraint encodes the condition: *affinity*(*m, x*) ∧ *x* ≥ *θ*. We ensure updates to the background knowledge performed by the `PyLMLF` implementation of the LMLF procedure ensure that *θ* values either stay the same or increase on each iteration. This ensures the constraint-generalisation condition is not violated. In the description of the experimental method below, we will denote the trivial case of not having any constraints as 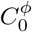; the case with only target-agnostic constraints as 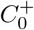; and the case with both target-agnostic and target-specific constraints as 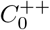.

#### Query

A query in our experiments is a restricted (unambiguous) English statement: *Generate valid molecules that do not belong to any known database* The query, in combination with the available background knowledge and constraints, is assembled to construct the prompt for the LLMs.

#### Algorithms and Machines

All the experiments are conducted using a Linux (Ubuntu) based workstation with 64GB main of main memory, 16-core Intel Xeon 3.10GHz processors. All the implementations are in Python3, with API calls to the respective model engines for GPT-3.0 and PaLM. We use RDKit (version: 2022.9.5) for computing molecular properties and GNINA 1.0 for computing docking scores of molecules.

### 4.3 Method

Our method is straightforward:

1. For each biological target *T ∈* {*JAK*2, *DRD*2}:
  a. For each LLM *L ∈* {*GPT* 3, *PaLM*}
    i. Let *M*_*T,i*_ be the set of molecules returned by `PyLMLF` provided with LLM *L*, background *B*_0_ and constraints 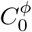 for target *T*;
    ii. Let 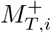 be the set of molecules returned by `PyLMLF` provided with LLM *L*, background *B*_0_ and constraints 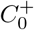 for target *T*; and
    iii. Let 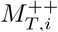 be the set of molecules returned by `PyLMLF` provided with LLM *L*, background *B*_0_ and constraints 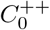 for target *T*.
    iv. Compare the sets *M*, 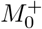 and *M* ^++^ quantitatively using the distribution of estimated target-specific binding affinities
    v. Compare the sets *M, M* ^+^ and *M* ^++^ qualitatively using assessments by domain-specialists

The following additional details are relevant:

- We make API calls to `text-davinci-003` for GPT-3.0 and `text-bison-001` for PaLM. For both LLMs, *temperature* is set to 0.7.
- The upper-bound on the number of iterations (*k* in Procedure 1) is 10.
- In our constraint *C*, we use a threshold of 7 on binding affinity for the first 5 iterations and 8 for the next 5 iterations.
- Quantitative comparison of performance is done as follows. For any set of molecules, we obtain a histogram of binding affinities, to act as an estimate of the probability distribution of affinities. Comparison of any 2 sets of molecules *M*_1_ and *M*_2_ is done by using the non-parametric Mann-Whitney U test on these estimated distributions. If *p* < 0.05 then we reject the null hypothesis that *M*_1_ and *M*_2_ are from the same distribution. If the null hypothesis is rejected, and the median values of binding affinities from *M*_1_ are higher than those from *M*_2_, then we will say the performance of the procedure generating *M*_1_ is better (respectively equal or worse).

### 4.4 Results

In the following, we use `GPTLF` to denote `PyLMLF` provided with GPT3, background *B*_0_, and constraints 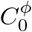; `GPTLF` ^+^ to denote `PyLMLF` provided with GPT3, background *B*_0_ and constraints 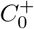; and `GPTLF` ^++^ to denote `PyLMLF` with GPT3, background *B*_0_, and constraints 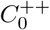. Similarly for `PaLMLF`, `PaLMLF` ^+^ and `PaLMLF` ^++^. Summaries of quantitative results are in Table 1. Histograms showing the distribution of estimated binding affinities is in Figure 2. It is evident from both the tabulation and diagrams that for both targets and both LLMs: (a) LLMs are capable of generating molecules without any constraints 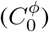; (b) When the LLMs are provided target-agnostic constraints 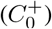, performance is better than without constraints (case (a) above); and (c) when the LLMs are provided with both target-agnostic and target-specific constraints 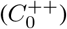, performance is better than with just generic constraints (case (b) above). These results provide quantitative support to the conjecture that logical feedback is beneficial in using LLMs to generate potential leads.

**Table 1:** Statistics of docking scores, computed using GNINA 1.0 for the GPT-3 and PaLM generated molecules against (Left) JAK2 and (Right) DRD2 as the drug-targets.

| Method | Mean | Median | S.D. | Range |
| --- | --- | --- | --- | --- |
| GPTLF | 6.50 | 6.61 | 1.07 | 4.21 – 7.55 |
| GPTLF<sup>+</sup> | 7.53 | 7.57 | 0.88 | 4.83 – 8.29 |
| GPTLF<sup>++</sup> | 7.74 | 7.71 | 0.30 | 7.18 – 8.52 |
| PaLMLF | 6.09 | 6.01 | 0.77 | 3.86 – 7.55 |
| PaLMLF<sup>+</sup> | 6.12 | 6.24 | 0.69 | 4.65 – 8.11 |
| PaLMLF<sup>++</sup> | 7.20 | 7.40 | 0.58 | 7.03 – 8.36 |
| GPTLF | 6.48 | 6.51 | 0.75 | 5.33 – 7.42 |
| GPTLF<sup>+</sup> | 7.32 | 7.21 | 0.32 | 6.45 – 8.02 |
| GPTLF<sup>++</sup> | 7.66 | 7.71 | 0.29 | 7.25 – 8.29 |
| PaLMLF | 6.05 | 6.13 | 1.05 | 4.33 – 7.49 |
| PaLMLF<sup>+</sup> | 6.22 | 6.23 | 0.48 | 5.67 – 8.24 |
| PaLMLF<sup>++</sup> | 7.60 | 7.55 | 0.37 | 7.01 – 8.19 |

**Figure 2.**
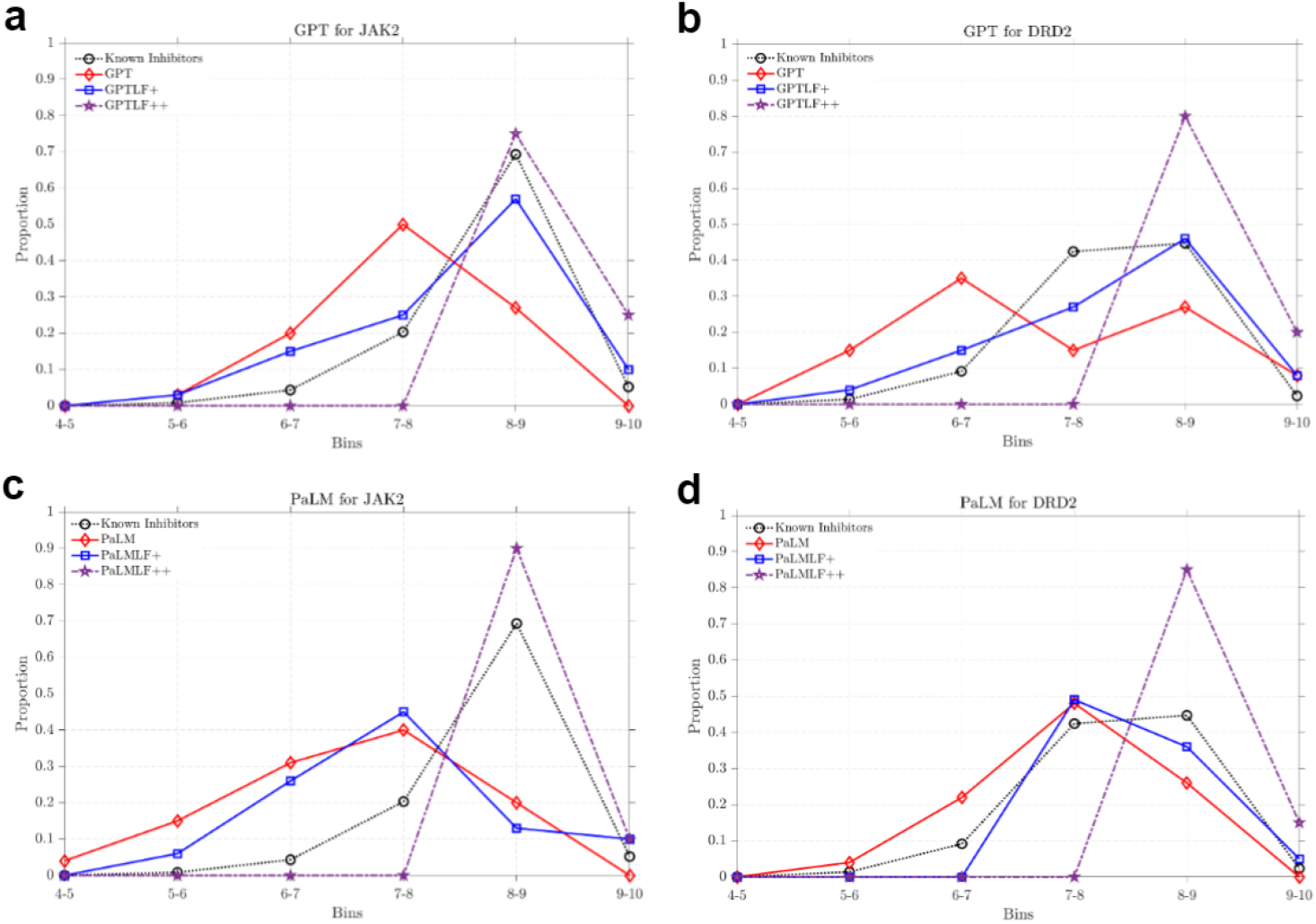
Plot showing the distribution of binding affinities for molecules generated by the LLMs and the two drug targets.

#### Qualitative Assessment by Chemists

The chemists were provided with 20 molecules each of potential JAK2 and DRD2 inhibitors (10 of each were generated by `GPTLF` ^++^ and `PaLM` ^++^, although the chemists were not told which LLM was involved). A summary of the assessment made by the chemists is reproduced below:

#### Efficacy

(a) *JAK2*. A set of 13 JAK2-selective functional groups were identified based on patent literature and used for exact substructure match against the generated molecules. From the search, 3 generated molecules (15%) were found to have at least one JAK2-selective functional group. 4,6-Diamino pyrimidine, morpholine, [1,2,4]-triazolo[1,5-a]pyridine and 3-amino pyrazole groups were predominantly observed to enhance JAK2 selectivity in the generated small molecules (b) *DRD2*. A set of 11 functional groups were identified from patented DRD2 inhibitors. Since these functional groups have been proven to enhance selectivity toward DRD2, an exact substructure match was performed with the generated small molecules using RDKit, to identify the presence of these selectivity features from patent literature. From the search, 8 generated molecules (40%) were found to have at least one DRD2-selective functional group indicating the model’s capability to optimize molecular features to capture selectivity, based on the docking score observed. Dimethyl piperazine and chlorobenzene was observed to be the predominant DRD2-selective group among the

#### Novelty

10 of the 11 molecules which contain JAK2 or DRD2 selective functional groups of interest also had less than 0.75 Tanimoto similarity to the existing JAK2 or DRD2 inhibitors, respectively. Therefore, it can be interpreted that although some fractions of the molecules generated have less similarity to existing inhibitors, the presence of selective functional groups indicates their potential to act as novel and selective inhibitors for the target protein of interest.

#### Overall

It appears that the model has learned to generate more inhibitors with better similarity to existing JAK2 inhibitors (50%) compared to DRD2 (15%). But this difference is compensated by the fact that 40% of generated DRD2 inhibitors have highly selective functional groups, while only 15% of generated JAK2 inhibitors have selective groups. Hence, the model exhibits a balance in generation of similar and novel molecules depending on the nature and diversity of the training dataset used for the target protein of interest.

We now report on some additional comparisons that do not directly impinge on the experimental conjecture in Sec. 4.1, but are nevertheless of interest to practitioners. First, the quantitative and qualitative assessments provide us with an opportunity to compare the capabilities of GPT3 and PaLM under controlled conditions. It is evident from the tabulation in Table 1 suggest the LMLF using GPT appears to be better than using PaLM. However, the expert assessment provides a slightly different story. Of the 11 molecules identified to be possibly effective JAK2 or DRD2 inhibitors, 7 were obtained using GPT3 and 4 were obtained using PaLM. Of the possible inhibitors that were also identified as being possibly novel, 7 were from GPT3 and 3 were from PaLM. These numbers are indicative of some benefit in using GPT. However the differences are not statistically significant.

Secondly, we are able to perform a comparison of the use of LLMs against baselines provided by: (a) known inhibitors of JAK2 and DRD2; and (b) results reported on the same dataset(s) on novel lead-generation. Results have been reported on the JAK2 dataset most recently in (Dash et al. 2021). This uses a combination of 2 variational autoencoders (VAEs) for generating molecules, and a graph-based neural network (GNN) that acts as a discriminator, which found to be perform better than previous reports (in (Krishnan et al. 2021)) using reinforcement learning in combination with an MLP. Table 2 compares and shows both LLMs perform better than the VAE-GNN combination. To some extent, this is unsurprising for 2 primary reasons: (1) the LLMs have access to substantially more information than the VAE-GNN model, and (2) the VAE-GNN model does not have access to the constraint(s) on the binding affinity of molecules. It is relatively straightforward to develop a variant of LMLF for other kinds generative models like the VAE-GNN model. This would address (2), but it is unclear how the gap in (1) can be bridged.

**Table 2:** Summary statistics of the distribution of estimated binding affinities for molecules generated by the LLMs with logical feedback, in comparison to those of known inhibitors, and molecules generated by the VAE-based generator (VAE -GNN) in (Dash et al. 2021). Left is for JAK2, and right is for DRD2 (‘-’ denotes ‘data not available’).

| Method | Mean | Median | S.D. | Range |
| --- | --- | --- | --- | --- |
| Known Inhibitors | 7.26 | 7.42 | 0.64 | 4.19 – 8.34 |
| VAE-GNN | 6.53 | 6.95 | 1.18 | 2.10 – 8.18 |
| GPTLF++ | 7.74 | 7.71 | 0.30 | 7.18 – 8.52 |
| PaLMLF++ | 7.20 | 7.40 | 0.58 | 7.03 – 8.36 |
| Known Inhibitors | 6.86 | 6.97 | 0.63 | 4.21 – 8.31 |
| VAE-GNN | - | - | - | - |
| GPTLF++ | 7.66 | 7.71 | 0.29 | 7.25 – 8.29 |
| PaLMLF++ | 7.60 | 7.55 | 0.37 | 7.01 – 8.19 |

## 5 Related Work

Over the last few years, deep generative models have been used quite successfully in generating novel compounds for specific biological targets and with desired molecular properties. A comprehensive review of some of the techniques can be found in (Sousa et al. 2021). Among these techniques, deep sequence models such as variational autoencoders (VAE), deep structure-based models such as graph neural networks (GNNs) are shown to be very effective. Some of these techniques are paired with each other and also with reinforcement learning to allow these models to be biased towards generating models with desired properties (Jin, Barzilay, and Jaakkola 2018; Gómez-Bombarelli et al. 2018; Liu et al. 2018; Krishnan et al. 2021; Cao and Kipf 2022). It has been shown also that deep sequence models can gain substantial advantage over classic deep models, mentioned above, by infusing them with some form of domain-knowledge such as logical rules, constraints, available facts (Dash et al. 2022). For instance, in (Dash et al. 2021) the authors show that VAEs are significantly better if provided with chemical knowledge via a Bottom-Graph Neural Network (Dash, Srinivasan, and Baskar 2022) for conditional molecule generation.

With the recent surge in development of LLMs, the field of AI for natural science is witnessing increasing adoption of LLMs in solving interesting problems such as molecular property prediction, molecule optimisation, compound discovery and the like (Castro Nascimento and Pimentel 2023; Kang and Kim 2023; Born and Manica 2022). LLMs with some (human) feedbaback are also adopted for molecular generation (Blanchard et al. 2023; Fang et al. 2023). Our present work is in a similar vein albeit with additional domain-knowledge and desired constraints used to progressively guide the LLM’s sampling engine to draw molecules from a more restricted joint distribution, allowing more novelty and diversity in molecule generation against a specific biological target.

## 6 Conclusion

In this paper, we have proposed a simple iterative procedure called LMLF that uses progressively alters the conditioning string provided to a large language model (LLM). The alterations are the result of testing answers generated by the LLM against domain-specific constraints represented in a formal, rather than natural language. On each iteration, LMLF requires the LLM to generate answers that satisfy the constraints, which are themselves automatically strengthened on subsequent iterations. We investigate the performance of LMLF in the area of lead-discovery, and find that the logical constraints, enforced using our proposed feedback mechanism, provide much more effective conditioning information to the LLM. As a result, we are able to use the internal knowledge contained within the LLMs much more effectively to generate potentially novel inhibitors for specific biological targets. We present quantitative results supporting this claim on two separate targets, using two different LLMs; and qualitative results in the form of preliminary assessments by computational chemists.

Large ‘foundation’ models that have been constructed with vast amounts of data can be seen as storehouses of factual and hypothesised knowledge that can be of great value in tackling complex tasks in areas like drug-design. But how are human problem-solvers–like chemists and biologists–to draw on such knowledge? A long recognised concern of mismatch between human- and machine-representations of knowledge suggests that this is not an easy task. On the surface, it would seem that this will not be a concern when using LLMs, given their capability to interact with humans in a natural language. However, this may not follow for at least two reasons. First, the issue is of a mismatch in representation (what concepts are being used), and not of communication language. For example, the machine may be using a concept for which there is no simple description in a natural language. We are not addressing this problem here. Secondly, the flexibility of natural language introduces ambiguity and imprecision. Thus, human-concepts can be conveyed to a machine in many ways, not all of which may map to the same machine-concepts (however mismatched). The latter issue poses difficulties in using LLMs in a controlled manner. The position adopted in this paper is that for certain kinds of scientific problems–like the lead-discovery tasks here–it is possible to side-step the second difficulty and still use LLMs effectively. Specifically, we are concerned with tasks for which we are able to formulate task-specific requirements in a sufficiently formal language, which can in turn be used in conjunction with an LLM. We suggest that this simple neuro-symbolic approach could provide an effective basis for using LLMs in closed-loop scientific discovery of the kind envisaged in (Zenil et al. 2023).

## Data and Code Availability

All the data and codes are available at https://github.com/Shreyas-Bhat/LMLF.

## Acknowledgements

RA and AS acknowledge the funding from DBT NNP (Grant No. BT/PR40236/BTIS/137/51/2022. The authors thank Shreyas V. for his assistance in getting the PaLM API running. AS is a TCS-affiliated Professor and the Head of the Anuradha and Prashanth Palakruti Centre for Artificial Intelligence Research (APPCAIR), BITS Pilani. He is also a Visiting Professor at the Centre for Health Informatics, Macquarie University, Sydney; and a Visiting Professorial Fellow at the School of CSE, University of New South Wales, Sydney. This work was initiated when TD was affiliated with BITS Pilani, Goa Campus. RA is a faculty member of APPCAIR, BITS Pilani, Goa Campus.

## Appendix

This is a supplementary document accompanying the main manuscript. Here we expand on some procedures and implementations used and provide additional results. We also provide a detailed analysis from two chemists of some novel target-specific molecules generated using LMLF.

### A The PyLMLF Implementation of LMLF

The implementation `PyLMLF` employs a form of informed sampling, in which *n* new molecules are drawn calling the LLM *n* times, each time providing the LLM with a possibly different molecule to act as a “seed” for the generator. This entails a second loop within the implementation, as shown below in Procedure A.2. The *AssemblePrompt* procedure is the statement in Line 7. If *Q* is the string `“Generate a molecule that is valid and not in any known database”` the prompt computed on Line 7 will be a string like `“Generate a molecule that is valid and not in any known database and similar to 1 m”`, where `m` is some text-based representation of the molecule (in this paper, these are SMILES strings). i The steps in Line 12–13 implement *UpdateBack*; and the step in Line 14 implements *GeneraliseConstraint*.

#### Procedure A.2: The PyLMLF Implementation of LMLF

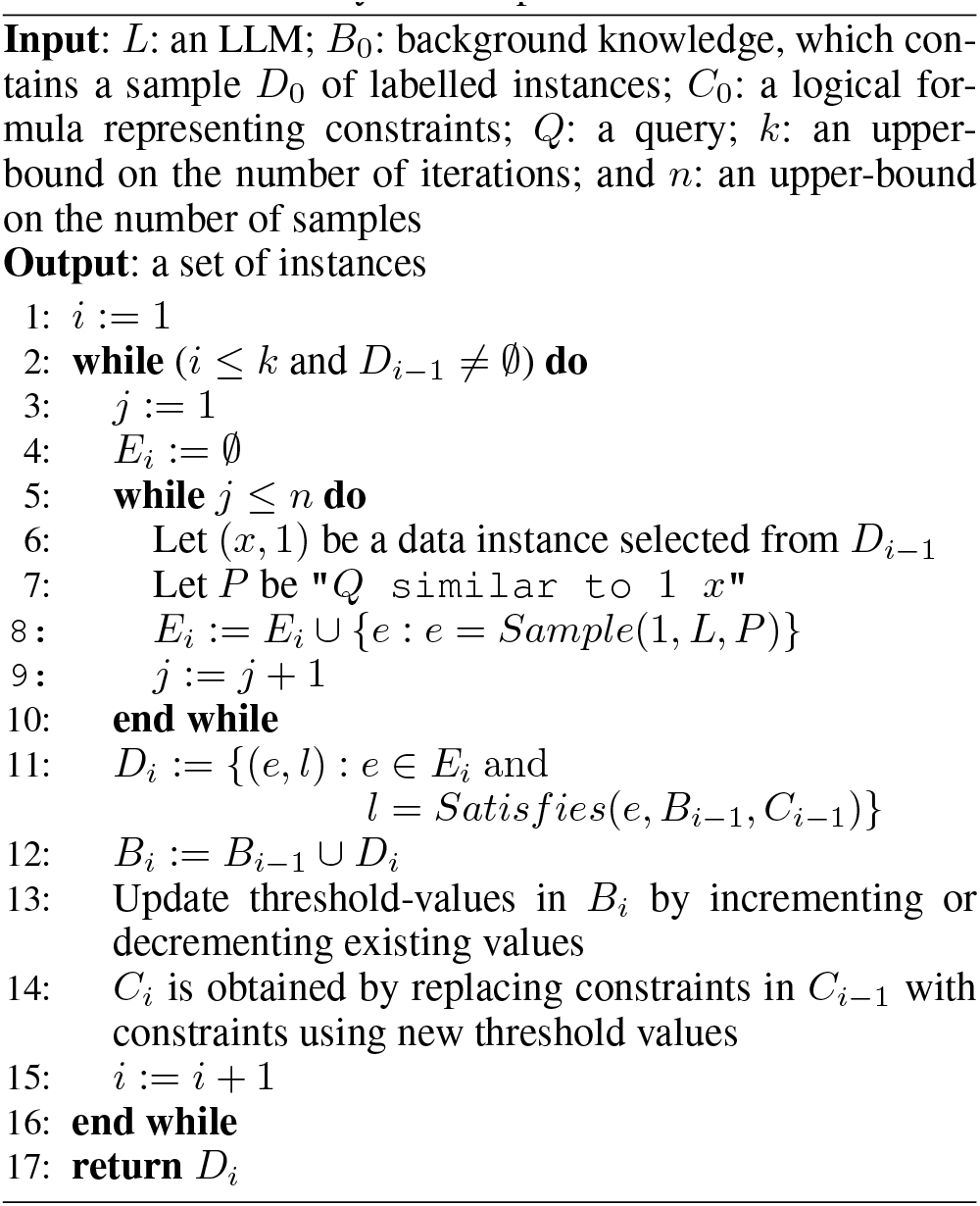

### B Additional Quantitative Results of LMLF-generated Molecules

In Fig. B.3–B.6 we provide some histogram-based summary of binding affinities of the molecules generated by the two LLMs for the two drug targets (JAK2 and DRD2). The results here clearly show that LLMs are indeed improving with the inclusion of the background knowledge and the logical feedback imposed via constraints.

### C Detailed Qualitative Analysis of LMLF-generated Molecules

Two computational chemists were provided with 20 generated molecules for each drug-target. Of these 20 molecules, they were not told which molecules were generated from which LLM (GPT3 and PaLM) to get their unbiased analysis and opinion.

#### C.1 DRD2 – 20 molecules

Upon comparison with 1798 existing DRD2 inhibitors curated from the ChEMBL database, 3 molecules (15%) were found to have high similarity (Tanimoto coefficient above 0.75) to known inhibitors (Fig. C.7).

To also understand the ability of the model to learn and incorporate functional groups necessary for better selectivity of the small molecules toward the DRD2 active site over its isoforms and homologs, a set of 11 functional groups were identified from patented DRD2 inhibitors. Since these functional groups have been proven to enhance selectivity toward DRD2, an exact substructure match was performed with the generated small molecules using RDKit, to identify the presence of these selectivity features from patent literature. From the search, 8 generated molecules (40%) were found to have at least one DRD2-selective functional group indicating the model’s capability to optimize molecular features to capture selectivity, based on the docking score observed. Dimethyl piperazine and chlorobenzene were observed to be the predominant DRD2-selective groups among the generated molecules (Fig. C.8).

#### C.2 JAK2 – 20 molecules

Upon comparison with 1103 existing JAK2 inhibitors curated from the ChEMBL database, 10 molecules (50%) were found to have high similarity (Tanimoto coefficient above 0.75) to known inhibitors (Fig. C.9).

Similar to the DRD2 case study, a set of 13 JAK2-selective functional groups were identified based on patent literature and used for exact substructure match against the generated molecules. From the search, 3 generated molecules (15%) were found to have at least one JAK2-selective functional group. 4,6-Diamino pyrimidine, morpholine, [1,2,4]-triazolo[1,5-a]pyridine and 3-amino pyrazole groups were predominantly observed to enhance JAK2 selectivity in the generated small molecules (Fig. C.10).

#### C.3 Comments on Novelty

From the results, it could be observed that more than 95% of the molecules which contain JAK2 or DRD2 selective functional groups of interest also had less than 0.75 Tanimoto similarity to the existing JAK2 or DRD2 inhibitors, respectively. Therefore, it can be interpreted that although some fraction of the molecules generated have less similarity to existing inhibitors, the presence of selective functional groups is indicative of their potential to act as novel and selective inhibitors for the target protein of interest.

Also, it appears that the model has learned to generate more inhibitors with better similarity to existing JAK2 inhibitors (50%) in comparison to DRD2 (15%). But this difference is compensated by the fact that 40% of generated DRD2 inhibitors have highly selective functional groups, while only 15% of generated JAK2 inhibitors have selective groups. Hence, the model exhibits a balance in the generation of similar and novel molecules depending on the nature and diversity of the training dataset used for the target protein of interest.

**Figure B.3:**
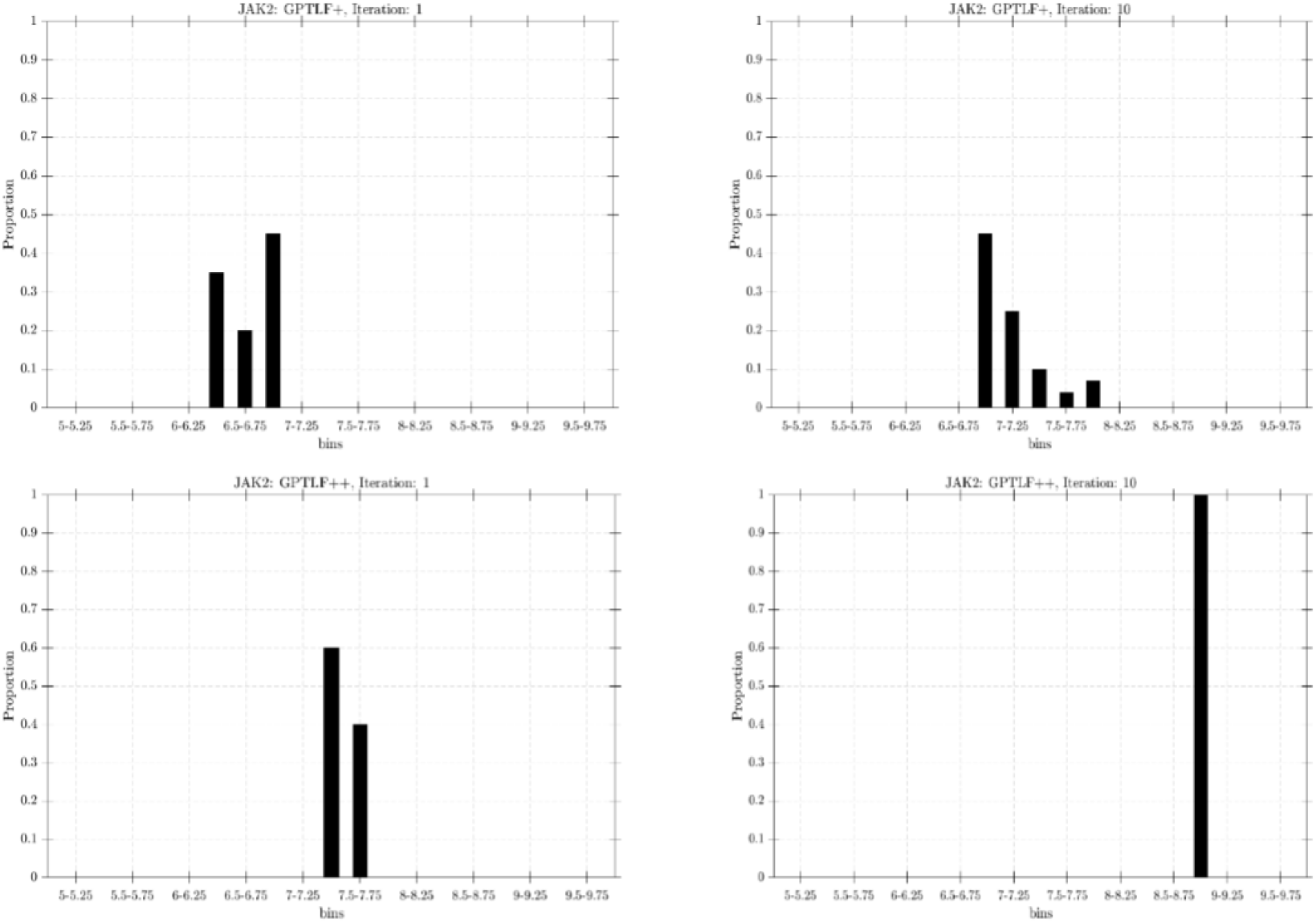
Histogram of binding affinities for the molecule generated by `GPTLF` ^+^ (top) and `GPTLF` ^++^ (bottom) for JAK2 in the first and last iterations.

**Figure B.4:**
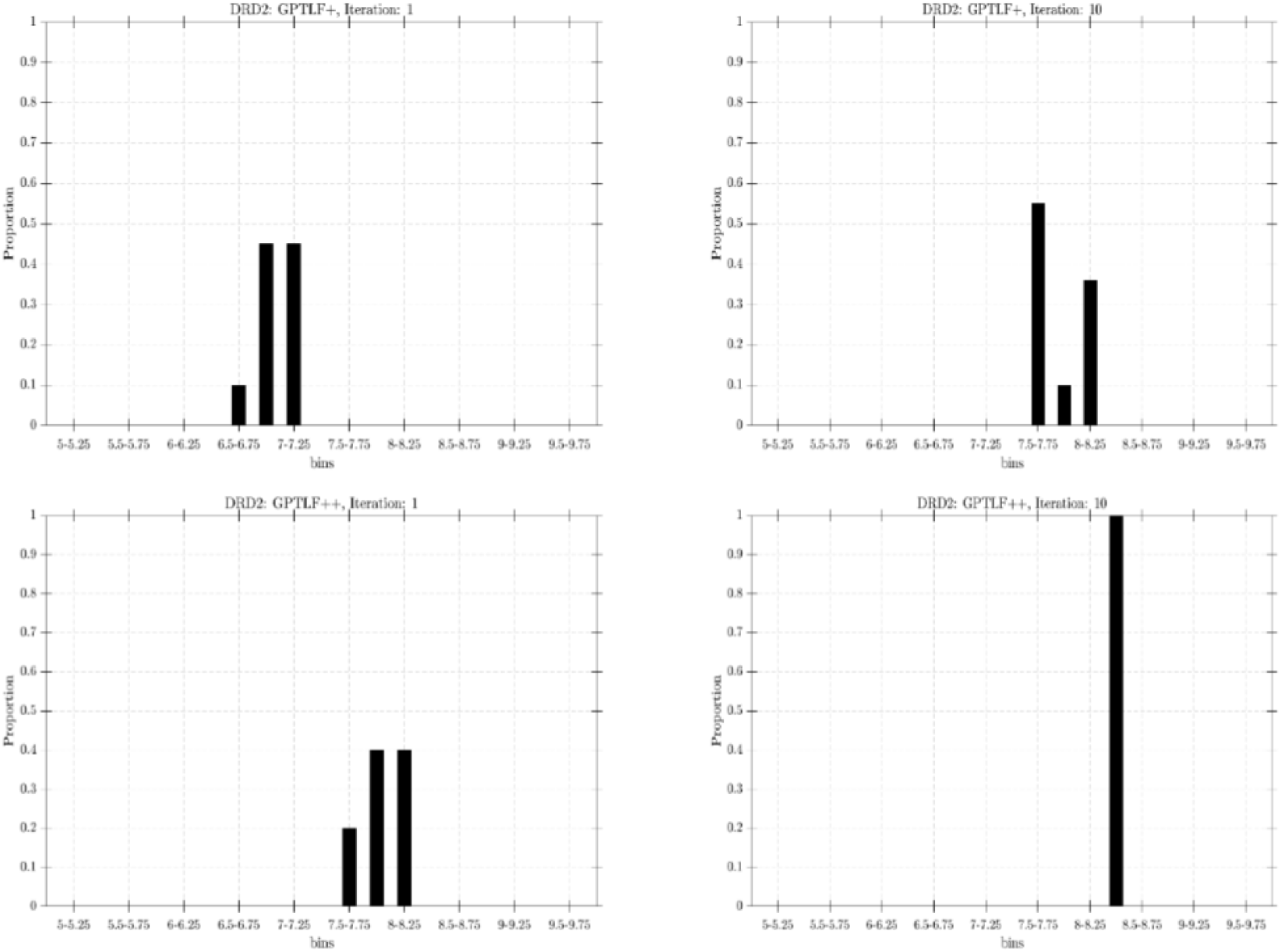
Histogram of binding affinities for the molecule generated by `GPTLF` ^+^ (top) and `GPTLF` ^++^ (bottom) for DRD2 in the first and last iterations.

**Figure B.5:**
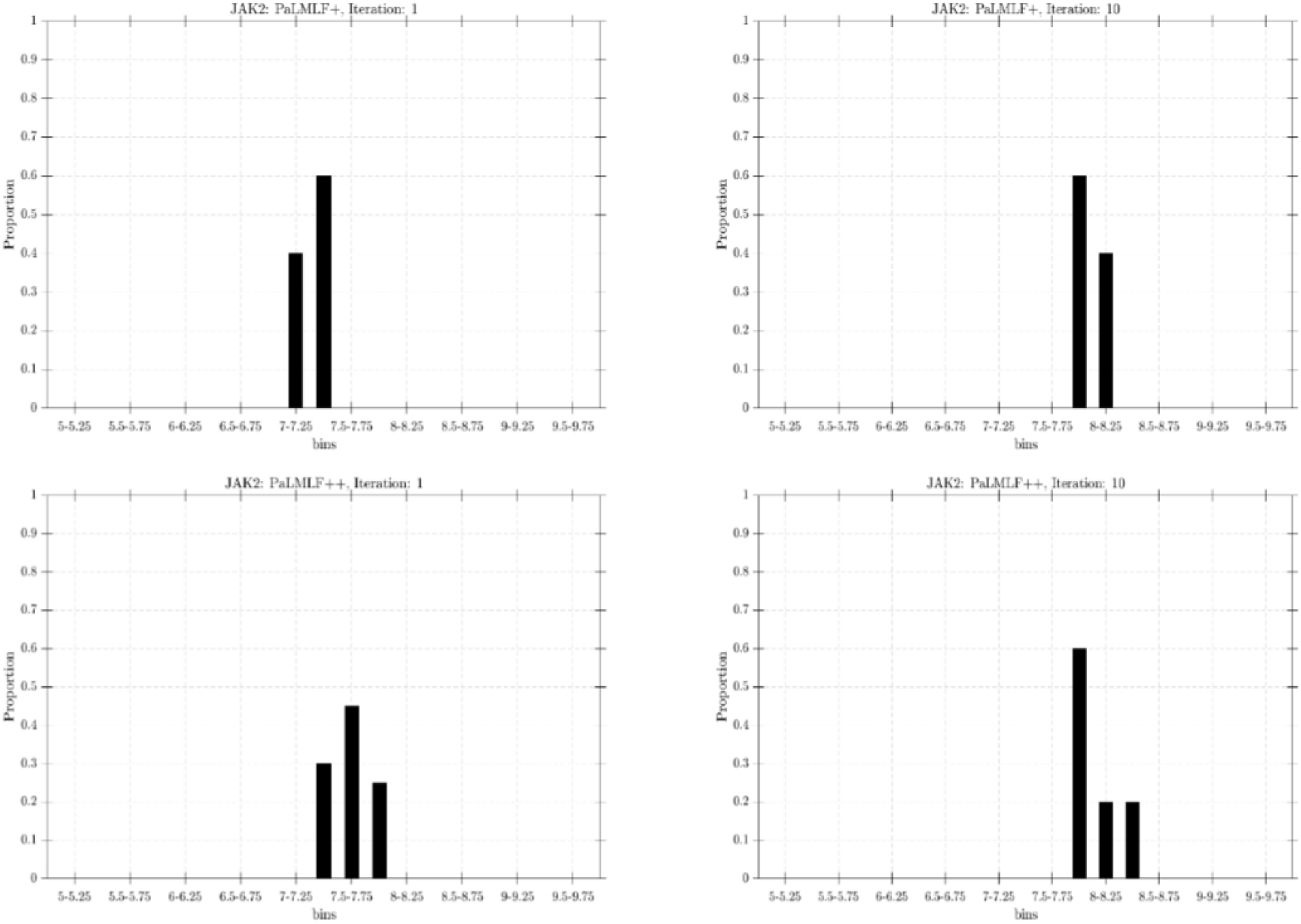
Histogram of binding affinities for the molecule generated by `PaLMLF` ^+^ (top) and `PaLMLF` ^++^ (bottom) for JAK2 in the first and last iterations.

**Figure B.6:**
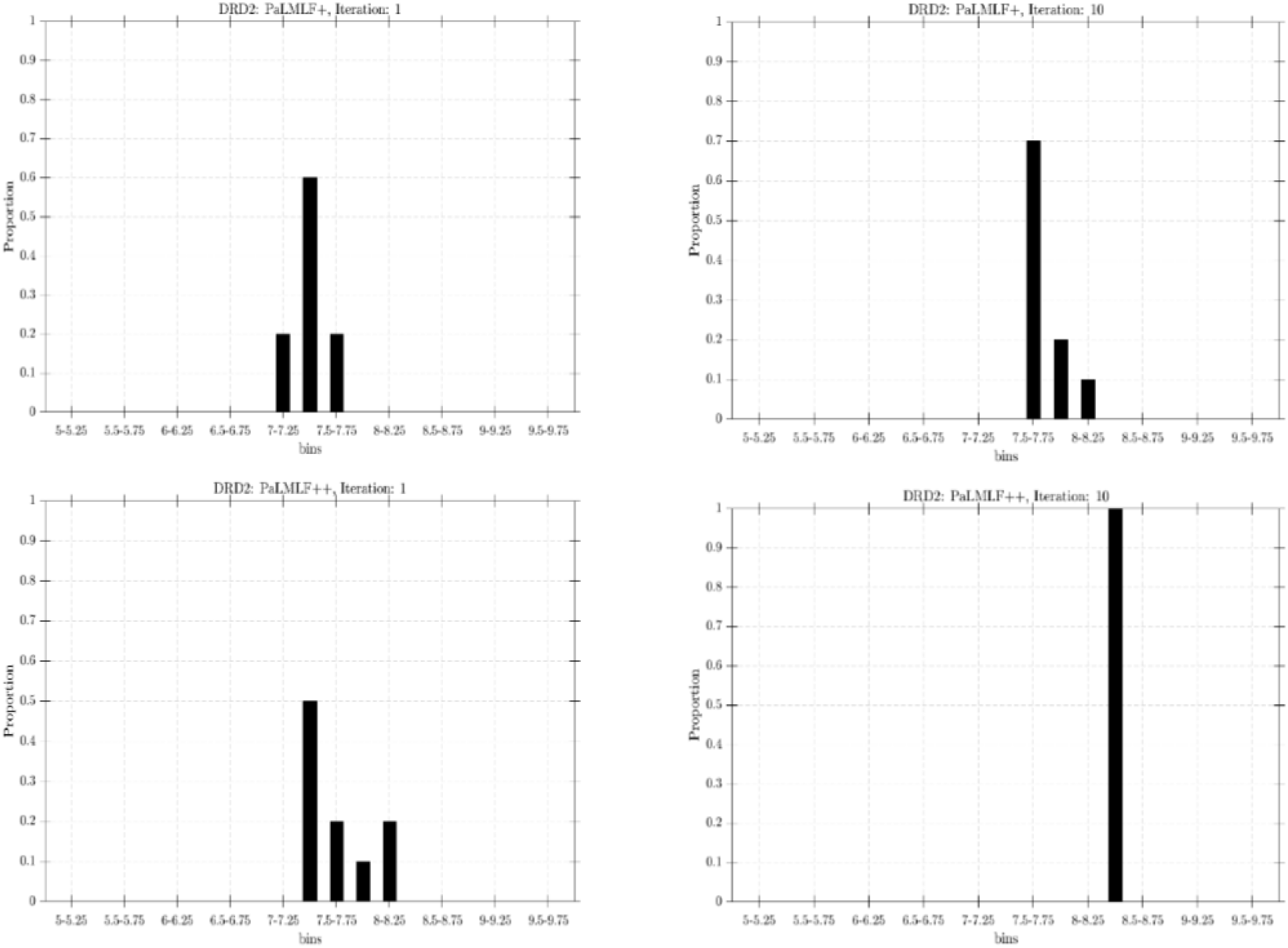
Histogram of binding affinities for the molecule generated by `PaLMLF` ^+^ (top) and `PaLMLF` ^++^ (bottom) for DRD2 in the first and last iterations.

**Figure C.7:**
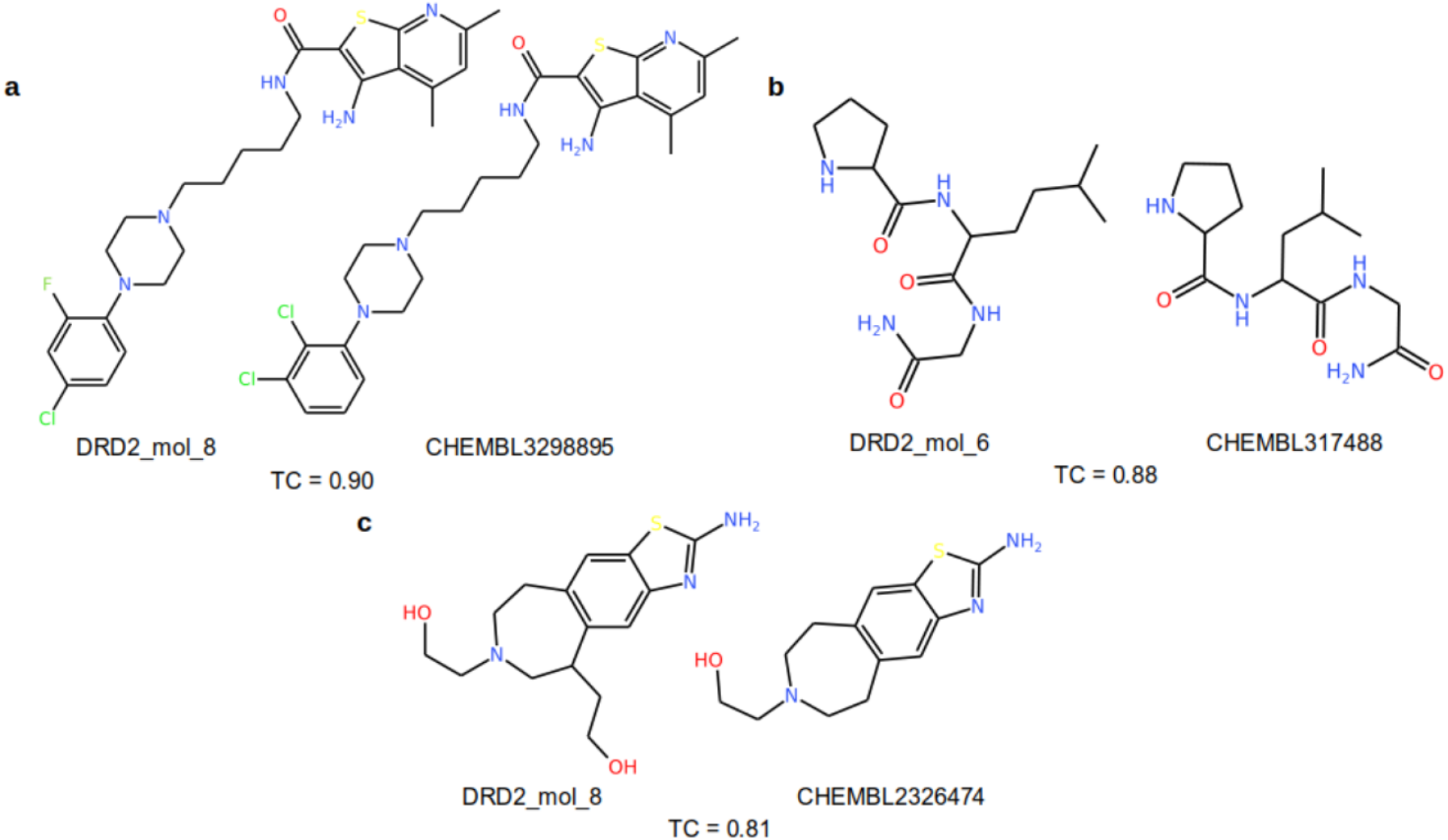
Examples of generated DRD2-specific small molecules with high similarity to existing DRD2 inhibitors. The ChEMBL ID of the known inhibitor is provided along with the similarity quantified in terms of the Tanimoto coefficient (TC).

**Figure C.8:**
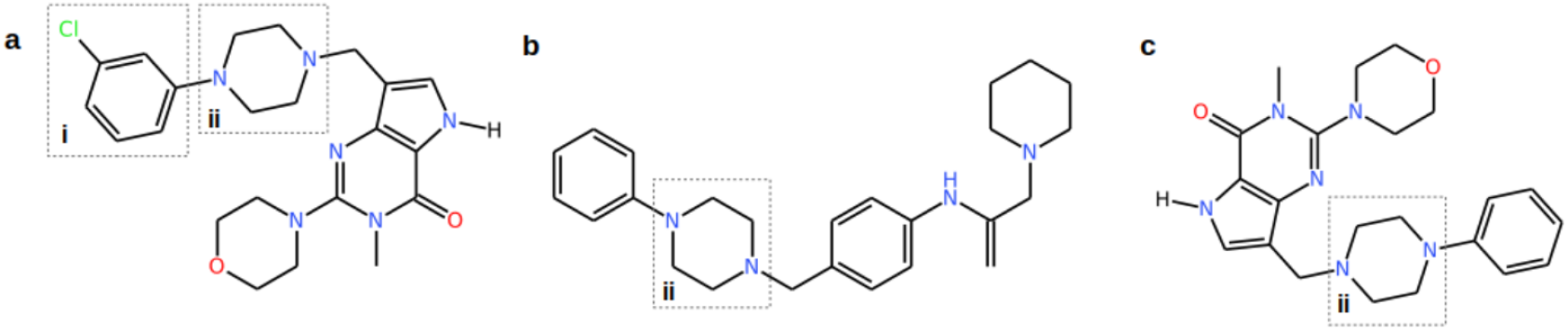
Examples of generated small molecules containing DRD2-selective functional groups observed in patent literature: (i) chlorobenzene and (ii) dimethyl piperazine.

**Figure C.9:**
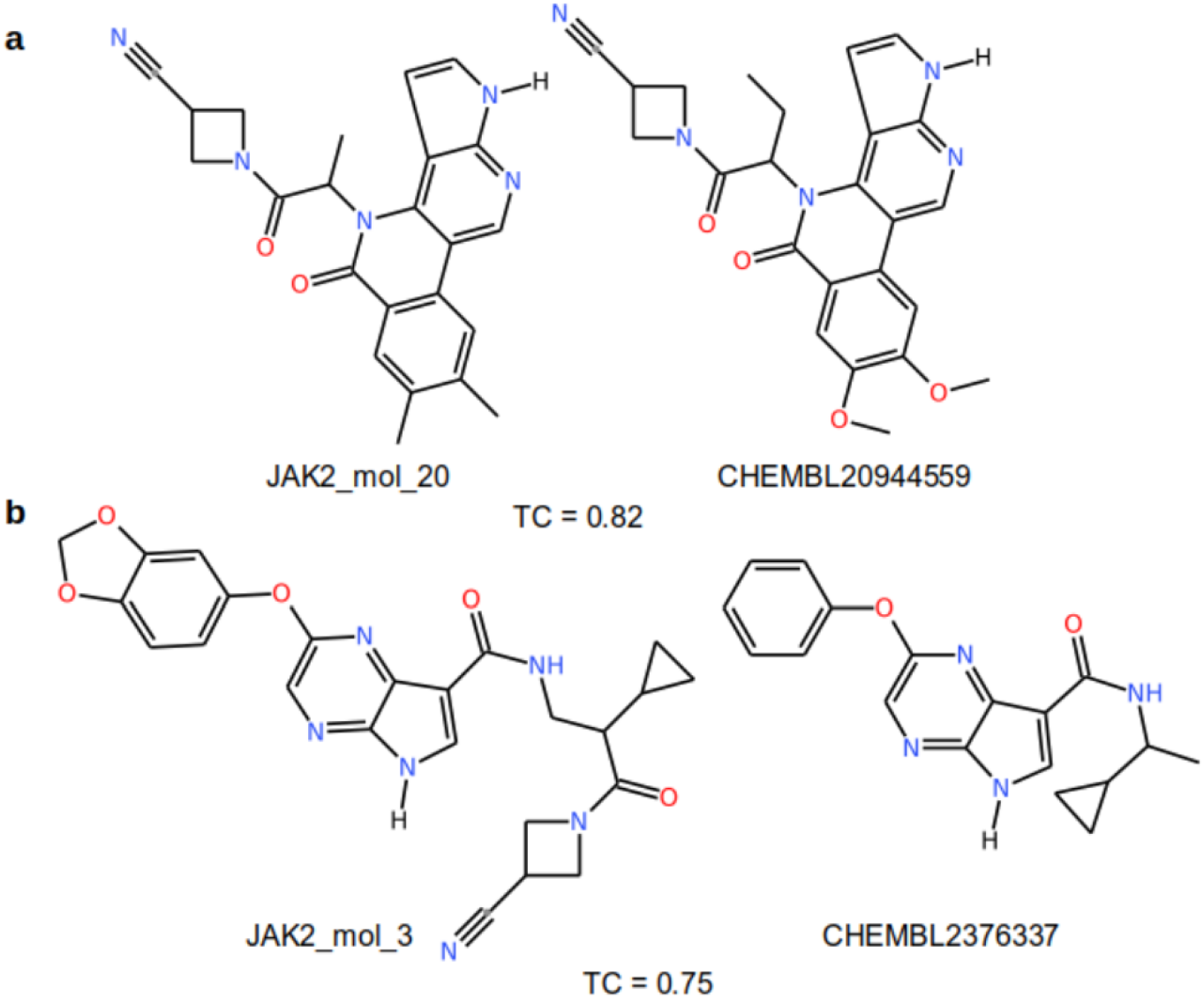
Examples of generated JAK2-specific small molecules with high similarity to existing JAK2 inhibitors. The ChEMBL ID of the known inhibitor is provided along with the similarity quantified in terms of the Tanimoto coefficient (TC).

**Figure C.10:**
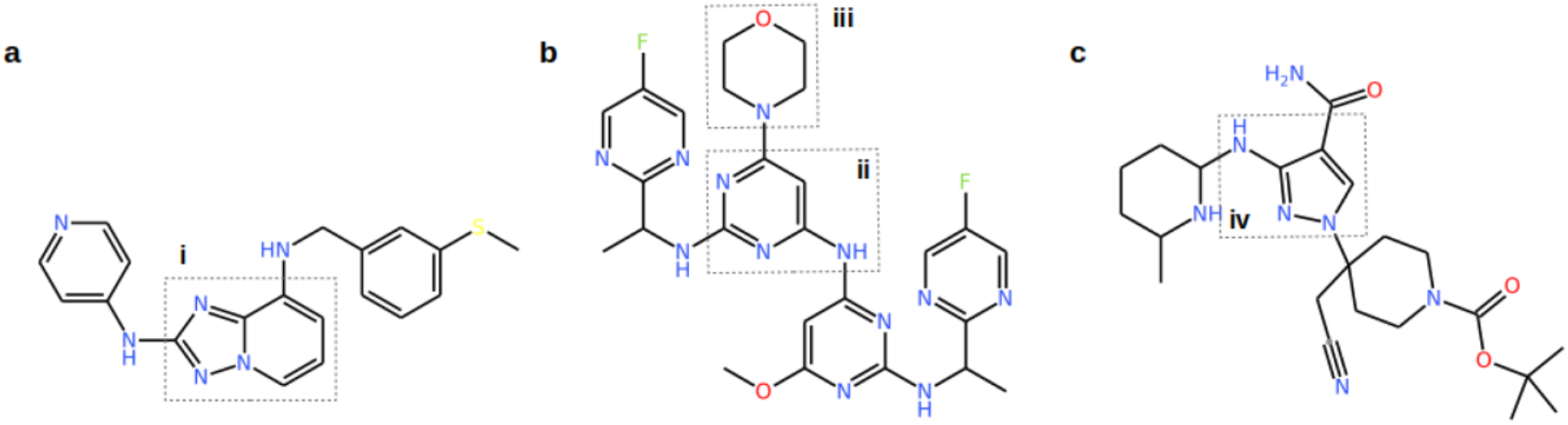
Examples of generated small molecules containing JAK2-selective functional groups observed in patent literature: (i) [1,2,4]-triazolo[1,5-a]pyridine, (ii) 4,6-diamino pyrimidine, (iii) morpholine and (iv) 3-aminopyrazole.

## Footnotes

1 Kopec distinguishes between ‘surface-level’ and ‘structural-level’ mismatches. The use of natural language addresses the surface-level mismatch, but it does not necessarily alleviate the mismatch between the concepts employed.

## Notes

https://github.com/Shreyas-Bhat/LMLF

https://www.ebi.ac.uk/chembl/

### Competing Interest Statement

The authors have declared no competing interest.

